# Bacterial killing by complement requires direct anchoring of Membrane Attack Complex precursor C5b-7

**DOI:** 10.1101/2019.12.17.877639

**Authors:** Dennis J Doorduijn, Bart W Bardoel, Dani AC Heesterbeek, Maartje Ruyken, Georgina Benn, Edward S Parsons, Bart Hoogenboom, Suzan HM Rooijakkers

## Abstract

An important effector function of the human complement system is to directly kill Gram-negative bacteria via Membrane Attack Complex (MAC) pores. MAC pores are assembled when surface-bound convertase enzymes convert C5 into C5b, which together with C6, C7, C8 and multiple copies of C9 forms a transmembrane pore that damages the bacterial cell envelope. Recently, we found that bacterial killing by MAC pores requires local conversion of C5 by surface-bound convertases. In this study we aimed to understand why local assembly of MAC pores is essential for bacterial killing. Here, we show that rapid interaction of C7 with C5b6 is required to form bactericidal MAC pores. Binding experiments with fluorescently labelled C6 show that C7 prevents release of C5b6 from the bacterial surface. Moreover, trypsin shaving experiments and atomic force microscopy revealed that this rapid interaction between C7 and C5b6 is crucial to efficiently anchor C5b-7 to the bacterial cell envelope and form complete MAC pores. Using complement-resistant clinical *E. coli* strains, we show that bacterial pathogens can prevent complement-dependent killing by interfering with the anchoring of C5b-7. While C5 convertase assembly was unaffected, these resistant strains blocked efficient anchoring of C5b-7 and thus prevented stable insertion of MAC pores into the bacterial cell envelope. Altogether, these findings provide basic molecular insights into how bactericidal MAC pores are assembled and how bacteria evade MAC-dependent killing.

## Introduction

When bacteria invade the human body, they are attacked by the immune system that tries to clear the infection. Gram-negative bacteria can be directly killed by the complement system, a family of proteins in blood and bodily fluids, that forms toroid-shaped Membrane Attack Complex (MAC) pores into bacterial membranes (1–4). MAC assembly is initiated when recognition molecules, such as antibodies and lectins, bind to bacterial surface structures and recruit early complement components to the target cell surface. This triggers activation of the complement system, a proteolytic cascade that labels the surface with convertase enzymes that cleave the central components C3 and C5 (5). Conversion of C5 into C5b, by C5 convertases, is a critical first step for the unidirectional step-wise assembly of the MAC pore (Fig. 1A). Upon formation of C5b, C6 rapidly binds metastable C5b to form a stable C5b6 complex (6). Subsequent association of C7 converts the hydrophilic C5b6 complex into lipophilic C5b-7 that uses an exposed membrane-binding domain of C7 to anchor to the target membrane (7,8). Membrane-bound C5b-7 subsequently recruits one copy of C8 and up to 18 copies of C9, which oligomerize and form of a 1.2 MDa transmembrane MAC pore (C5b-9) (9–11).

**Figure 1.**
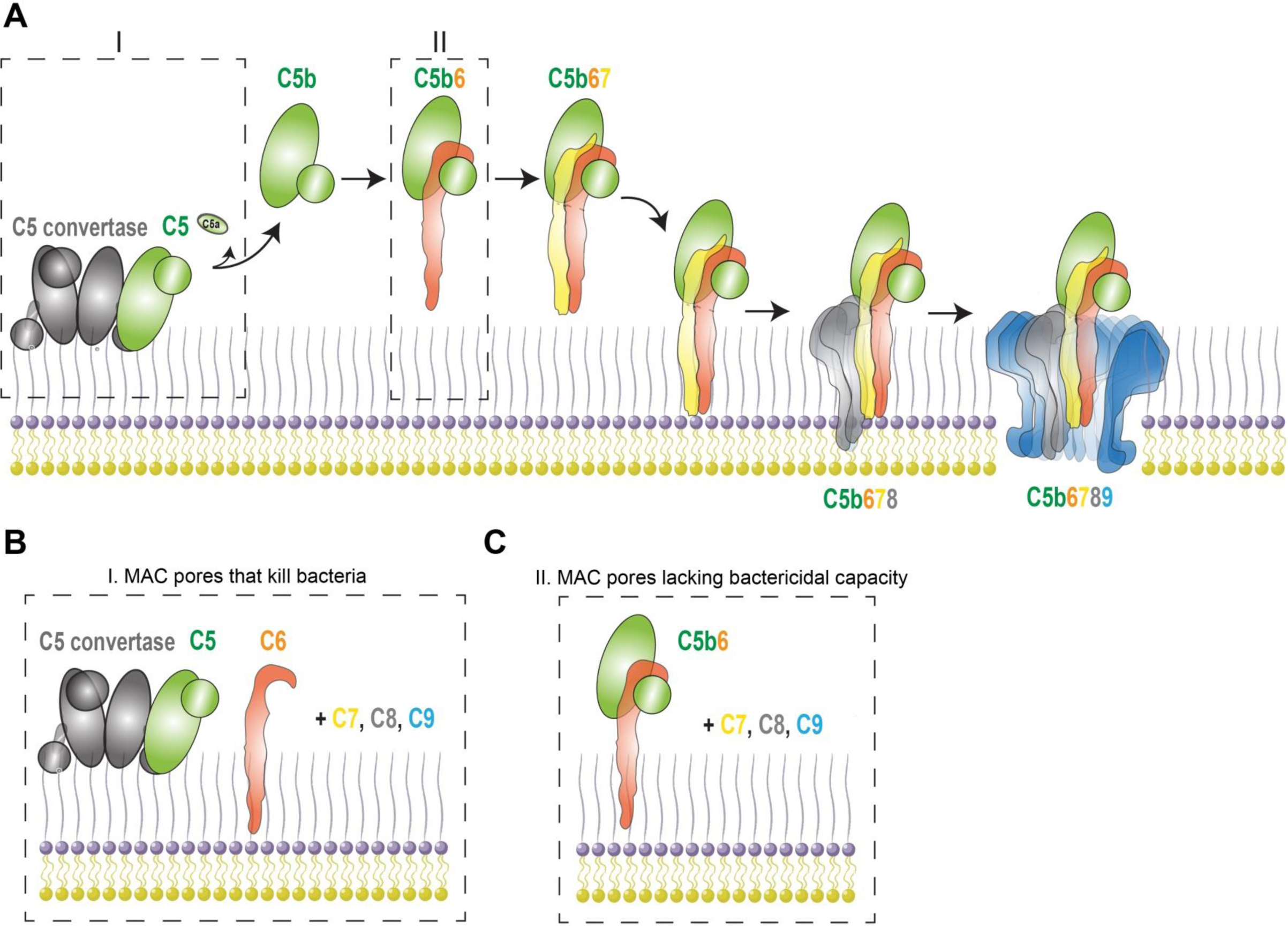
Step-wise assembly of the Membrane Attack Complex. (A) Illustration depicting how a C5 convertase bound to a bacterial surface triggers the assembly of the Membrane Attack Complex (MAC). The C5 convertase converts C5 into C5a and C5b. C5b immediately binds C6 which forms the bimolecular C5b6 complex. C5b6 binds to C7, which triggers conformational changes that exposes a lipophilic domain in C7 that anchors to membranes. C5b-7 then recruits C8, which causes initial membrane protrusion. C5b-8 then recruits multiple copies of C9 which forms a mebranolytic pore called the MAC. (B) Local C5 conversion and MAC assembly by a C5 convertase are essential to form a MAC pore that kills bacteria. (C) Isolated C5b6 can still trigger the assembly of MAC pores, but these MAC pores fail to kill bacteria.

Although the molecular steps of MAC assembly have been extensively studied on erythrocytes and model membranes (8,9,12–15), the mechanisms by which complement forms lytic MAC pores in the complex cell envelope of Gram-negative bacteria are less well-understood. Recently we studied how MAC pores cause perturbations to the Gram-negative outer membrane (OM) and inner membrane (IM) (16,17). Our data suggested that the proteolytic cascade preceding MAC assembly is crucial for the formation of bactericidal MAC pores that damage the bacterial IM. Whereas MAC pores formed from isolated C5b6 complexes effectively lysed single membrane models and erythrocytes, such complexes failed to kill Gram-negative bacteria (Fig. 1C). For MAC to kill Gram-negative bacteria, pore formation should be triggered via the conversion of C5 by surface-bound C5 convertases and subsequent local formation of C5b6 (Fig. 1B). Altogether this revealed that C5b6 can lose its capacity to form bactericidal MAC pores and that local assembly of C5b6 is a prerequisite to efficiently insert MAC pores into the bacterial OM and next damage the IM.

In this paper, we aimed to get more insight into MAC assembly on bacteria and uncovered that that rapid interaction with C7 is crucial for maintaining the bactericidal capacity of locally formed C5b6 on different complement-sensitive *E. coli* strains. Mechanistic studies show that C7 prevents release of C5b6 from the bacterial surface and thereby allows efficient anchoring of C5b-7 and subsequent insertion of MAC pores into the bacterial cell envelope. Additionally, we identify ‘complement-resistant’ *E. coli* strains that specifically prevent efficient anchoring of C5b-7 to the bacterial cell envelope. This highlights a novel mechanism how bacteria evade complement-dependent killing.

## Results

### Killing of *E. coli* by MAC pores requires rapid interaction between C7 and C5b6

To study different steps of MAC assembly on bacteria, we used our previously established assay to form MAC pores from purified complement components (16). *E. coli* bacteria were first labelled with C5 convertases through incubation in C5-depleted serum and then exposed to purified MAC components (C5, C6, C7, C8 and C9). As a read-out for bacterial killing, we measured IM damage using flow cytometry by the influx of a DNA dye (Sytox blue) that cannot pass an intact bacterial IM. As described previously, incubation of convertase-labelled bacteria with all MAC components simultaneously results in efficient IM damage in ~ 100% of bacteria for MG1655 (Fig. 2B, representative FACS plots shown in **S1)**.

**Figure 2.**
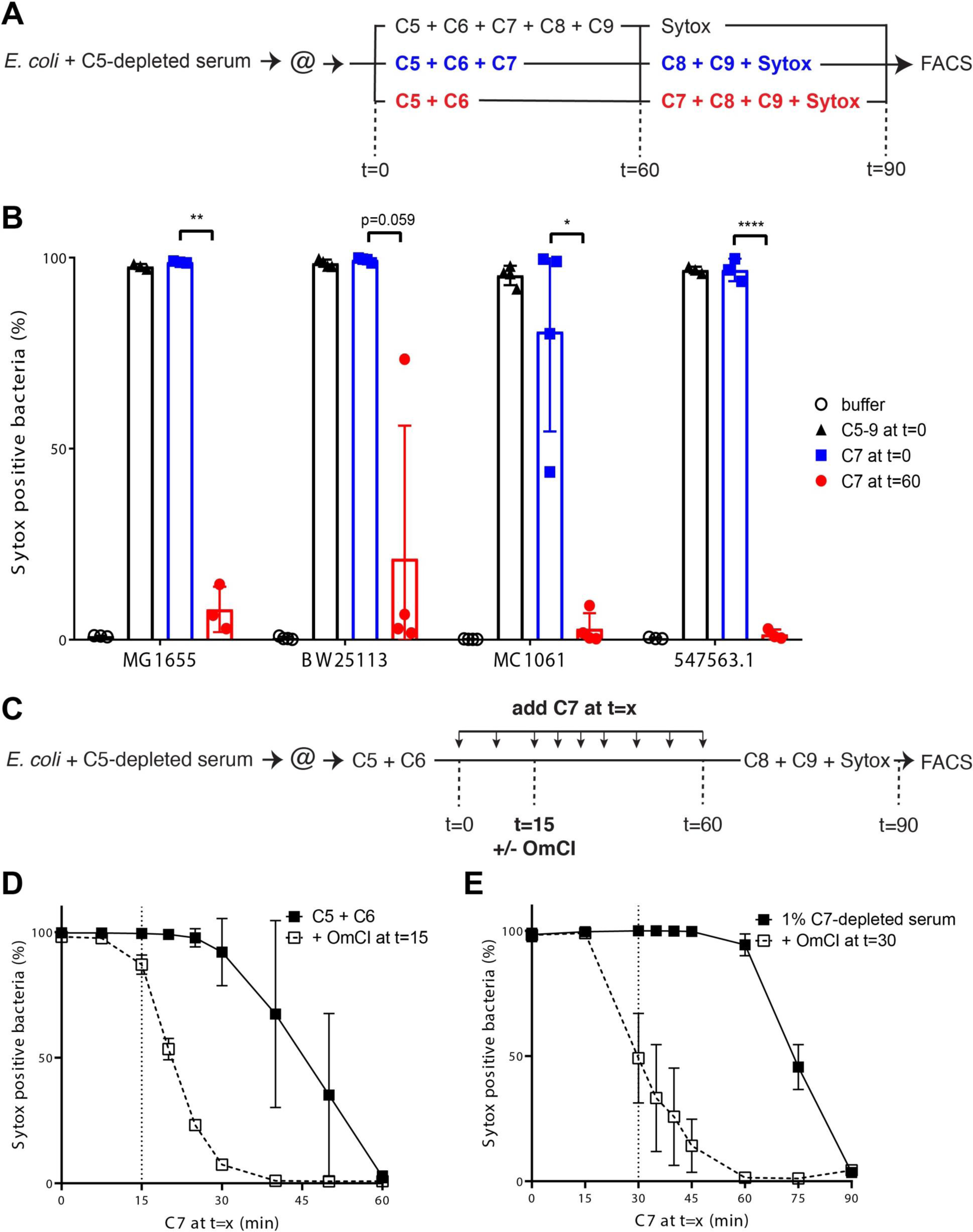
Killing of *E. coli* by MAC pores requires rapid interaction between C7 and C5b6. **(A)** Schematic overview of assay to determine the effect of the presence of C7 during C5b6 formation on bacterial killing by MAC pores. Bacteria were labelled with convertases in C5-depleted serum and washed (@) to get rid of remaining serum components. MAC pores were then either directly formed by adding all MAC components at t=0, by forming C5b6 in the presence of C7 at and adding C8 and C9 after 60 minutes or by forming C5b6 in the absence of C7 and adding C7, C8 and C9 after 60 minutes. Sytox is added to all bacteria after 60 minutes to analyze IM damage after 90 minutes (t=90) by flow cytometry. (**B)** The percentage of Sytox positive bacteria of four different complement-sensitive *E. coli* strains (MG1655, BW25113, MC1061 and 547563.1) which were treated as described in **(A)**. Samples with all MAC components at t=0 are represented in **black triangles**, samples with C5b6 formed in the presence of C7 in **blue squares** and samples with C5b6 formed in the absence of C7 in **red circles**. **(C)** Schematic overview of assay in **(D)** to determine the time-sensitivity of the interaction between C5b6 and C7 on bacterial killing by MAC pores. Convertase-labelled bacteria were incubated with C5 and C6 at t=0 and C7 was added at different points in time. At t=15, C5b6 formation was stopped by adding 25 µg/ml C5 conversion inhibitor OmCI. After 60 minutes, C8, C9 and Sytox were added to measure the percentage of Sytox positive cells by flow cytometry. **(D)** The percentage of Sytox positive bacteria of MG1655 bacteria which were treated as described in **(C)**. The solid line represents samples where C7 was added in time without stopping C5b6 formation, the dashed line represents samples where C5b6 formation was stopped with OmCI at t=15. (**E**) MG1655 was incubated with 1% C7-depleted serum followed by addition of C7 in time (solid line). To block C5b6 formation 25 µg/ml OmCI (dashed line) was added after 30 minutes. IM damage was determined by measuring the percentage of Sytox positive cells by flow cytometry after 90 minutes. Data represent mean +-SD (**B, D** and **E**) of at least 3 independent experiments. Statistical analysis was done using a paired 2-way ANOVA with Tukey’s multiple comparisons’ test in which all samples from one strain were compared with each other (**B**). Significance was shown as * p ≤ 0.05, ** p ≤ 0.01, *** p ≤ 0.001 or **** p ≤ 0.0001.

From earlier experiments on erythrocytes it is known that C5b requires an immediate interaction with C6 (6,12,18). However, the interaction of C5b6 complexes with C7, C8 and C9 is not known to be time-sensitive, because C5b6 complexes can be isolated and at a later moment mixed with C7, C8 and C9 to form pores on model membranes or erythrocytes (15,19–21). On Gram-negative bacteria however, we recently observed that C5b6, formed locally by surface-bound C5 convertases, loses its bactericidal potential by performing a washing step before adding C7 (16). This made us wonder whether the interaction of C7 with C5b6 is time-sensitive for bacterial killing. Here, we assessed how the presence or the absence of C7 during C5b6 generation affects the bactericidal capacity of C5b6. Instead of washing the bacteria before adding C7, we either added C7 at the beginning of the experiment or after 60 minutes of C5b6 generation. Bacterial killing was finally measured after C8 and C9 were added (schematic overview in Fig. 2A). C5b6 that was generated in the presence of C7, with C8 and C9 added after 60 minutes, efficiently damaged the IM of ~100% of bacteria for MG1655 (Fig. 2B, **S1**). This revealed that the presence of C8 and C9 during C5b6 generation was not required for C5b6 to retain its ability to trigger the formation of bactericidal MAC pores. However, if C7 was added 60 minutes after C5b6 generation was initiated, subsequent addition of C8 and C9 resulted in IM damage in ~6% of bacteria. (Fig. 2B, **S1**). Similar to these data on MG1655, we observed that MAC-mediated killing of other complement-sensitive *E. coli* strains (clinical isolate 547563.1 and laboratory strains BW25113 and MC1061) also required C7 to be present during C5b6 formation (Fig. 2B). In summary, we show that C5b6 requires a rapid interaction with C7 to maintain its bactericidal capacity.

To get more insight into how fast the interaction between C7 and newly generated C5b6 at the bacterial surface has to occur, C7 was added in a time-dependent manner during the generation of C5b6 on *E. coli* strain MG1655 (schematic overview in Fig. 2C). IM damage started to decrease when C7 was added 30 minutes after C5b6 generation was initiated and was completely lost after 60 minutes (Fig. 2D). Since C5 conversion, and thus C5b6 generation, is still ongoing during this assay, a C5 conversion inhibitor (OmCI) was added after 15 minutes to ensure that no new C5b6 could be generated after C7 was added. This resulted in a direct decrease of IM damage when C7 was added after C5b6 generation was stopped and IM damage was completely lost when C7 was added more than 15 minutes after C5b6 generation was stopped (Fig. 2D). Finally, we wanted to exclude that the absence of components naturally present in serum causes C5b6 to be dependent on the rapid interaction with C7. Bacteria were therefore incubated in 1% C7-depleted serum, which allows the generation of C5b6 while C7 was added in a time-dependent manner. In this set-up, IM damage started to decrease when C7 was added after 60 minutes or immediately when C7 was added after C5b6 generation was stopped with OmCI (Fig. 2E). Altogether, these data highlight that there is a time-sensitive interaction between C7 and C5b6 at the surface, which is required for bacterial killing by MAC pores in a variety of complement-sensitive *E. coli* strains.

### C7 prevents release of C5b6 from the surface

We next wondered why rapid interaction between C7 and nascent C5b6 is crucial for bacterial killing by MAC pores. Since binding of C7 to C5b6 anchors C5b-7 to the membrane, we hypothesized that rapid interaction with C7 prevents release of C5b6 from the bacterial surface. Therefore, convertase-labelled *E. coli* were incubated with C5 and C6 in the absence (sup *E. coli* + C56) or presence of C7 (sup *E. coli* + C567) and bacterial supernatants and cell pellets were collected. Using Western blotting, we analyzed the presence of C5b, which can be detected by a shift in molecular weight of the α-chain (α) of C5 when C5a is cleaved (α’). In the absence of C7, C5b was found in the supernatant and was barely detectable in the cell pellet, whereas in the presence of C7, C5b was mainly detected in the cell pellet (Fig. 3A). Using a C5b6-specific ELISA (validated in **S2**), we found ten-fold more C5b6 complexes in the supernatant of bacteria where C5b6 was generated in the absence of C7 (Fig. 3B). These data suggest that C5b6 is more prone to release from the bacterial surface when C7 is not present.

**Figure 3.**
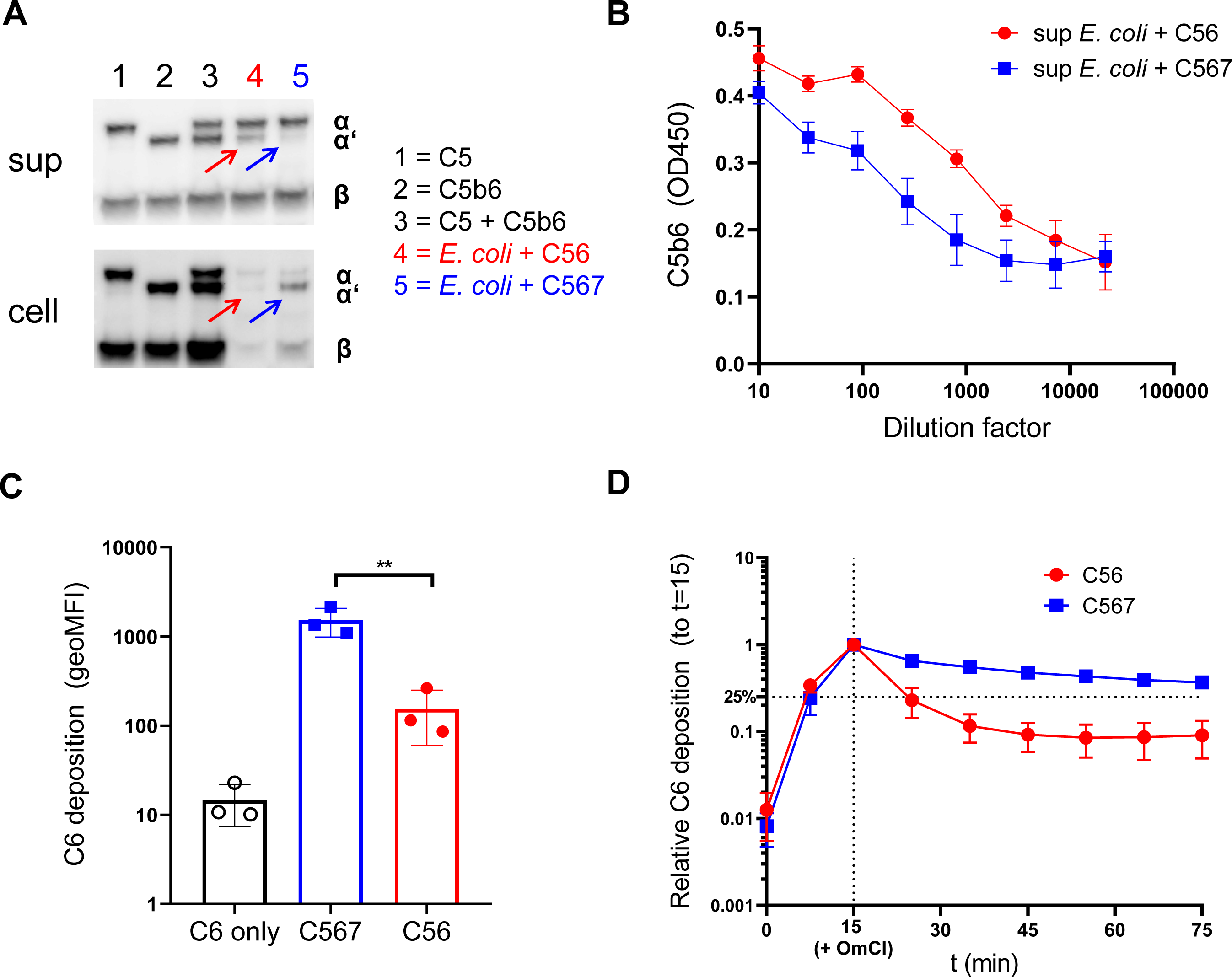
C7 prevents release of C5b6 from the surface. **A)** Representative Western blot for the detection of C5 and C5b in cell pellets and the supernatants taken from convertase-labelled *E. coli* MG1655 incubated with 100 nM C5, 100 nM C6 in the absence (**red arrows**) or presence (**blue arrows**) of 100 nM C7 for 60 minutes. As control, 50 pmol of C5, C5b6 or C5 + C5b6 was added to determine if the polyclonal anti-C5 binds both C5 and C5b to an equal extent. C5 is characterized by containing an uncleaved α-chain and β-chain, C5b by containing a cleaved α-chain (α’) and β-chain. **(B)** C5b6 ELISA of a titration of the supernatants used in **(A)**. C5b6 formed on bacteria in the absence of C7 is depicted in **red circles** and C5b6 formed in the presence of C7 in **blue squares**. A mixture of 10 nM C5 and 10 nM C6 was taken as control for specificity of the ELISA for C5b6. **(C)** Deposition of C6-Cy5 on convertase-labelled *E. coli* MG1655 was shown after 60 minutes with 10 nM C5, 10 nM C6-Cy5 in the absence or presence of 10 nM of C7. C6 deposition was measured by flow cytometry and plotted as the geometric mean of fluorescence intensity (geoMFI) of Cy5 of the bacterial population. (**D**) Relative deposition of C6-Cy5 in time for convertase-labelled bacteria incubated with C5, C6-Cy5 in the absence or presence of C7. At t=15, C5b6 formation was stopped by adding 25 µg/ml C5 conversion inhibitor OmCI (represented by the vertical dotted line). Relative C6 deposition was calculated by dividing the geoMFI at t=x by the geoMFI at t=15. Data represent mean +-SD (**B, C** and **D**) of at least 3 independent experiments (**A-D**). Statistical analysis was done using a paired one-way ANOVA with Tukey’s multiple comparisons’ test (**C**). Significance was shown as ** p ≤ 0.01.

To assess how fast C5b6 releases from the bacterial surface, we created fluorescently labelled C6. C6 was expressed with a C-terminal LPETG-tag and labelled with GGG-azide via a sortase enzyme (22), which allowed subsequent linking of C6-azide to DBCO-Cy5 via click chemistry. This labelling did not affect functionality of C6 (**S3**). C6 deposition on bacteria was observed in a C5-dependent manner and was ~ ten-fold higher when C5b6 was generated in the presence of C7 after 60 minutes (Fig. 3C). Next, convertase-labelled bacteria were incubated with C5 and C6 for 15 minutes, after which C5b6 generation was stopped with OmCI and the release of C5b6 was measured with time. In the absence of C7, C6 deposition decreased rapidly within 10 minutes to 25% of the signal at the moment when C5 conversion was stopped, and further decreased to ~8% of the signal after 60 minutes (Fig. 3D). By contrast, in the presence of C7, deposition of C6 was still at 70% of the initial signal 10 minutes after stopping C5 conversion and at 35% of the initial signal after 60 minutes (Fig. 3D). Altogether, these data indicate that C7 prevents release of C5b6 from the bacterial surface.

### C5b6 that is released from the bacterial surface loses its bactericidal potential

Since our data indicate that C7 is crucial to prevent the release of C5b6 from the surface, we hypothesize that C5b6 loses its bactericidal capacity upon release from the surface. To test this hypothesis, supernatant containing C5b6 that is released from the bacterial surface (sup *E. coli* + C56, used Fig. 3A) was tested for its bactericidal activity. The concentration of C5b6 in the supernatant was estimated at ~10 nM by ELISA (**S2**) and by an erythrocyte lysis assay (**S4**). Incubation of this supernatant containing released C5b6 together with C7, C8 and C9 did not result in IM damage on convertase-labelled MG1655 (Fig. 4A). By contrast, when uncleaved C5 and C6 was used together with C7, C8 and C9, the IM was efficiently damaged (Fig. 4A). C5b6 in the supernatant resulted in the deposition of fluorescently labelled C9 (Fig. 4B), but the deposition of C9 was approximately ten-fold lower compared to C5b6 that is generated at the bacterial surface. Even when the deposition of C9 was comparable to surface-generated C5b6, released C5b6 did not form MAC pores that damaged the bacterial IM. Altogether, these data indicate that C5b6 that is released from the bacterial surface loses its bactericidal potential.

**Figure 4.**
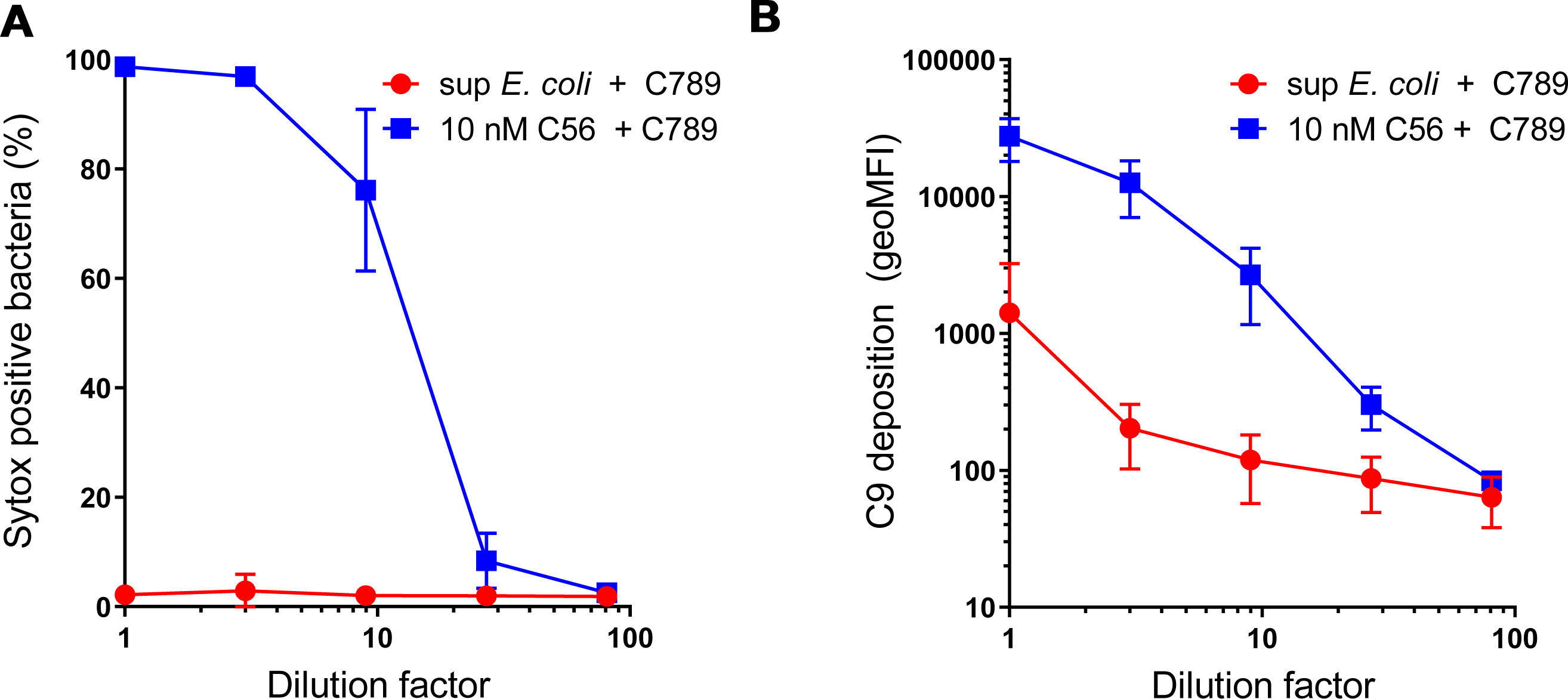
C5b6 that is released from the bacterial surface loses its bactericidal potential. Convertase-labelled *E. coli* MG1655 was treated with a titration of supernatant from *E. coli* + C56 described in Fig. 3A containing released C5b6 (**red circles**) or with 10 nM uncleaved C5 and 10nM C6 (**blue squares**) in the presence of 10 nM C7, 10 nM C8, 100 nM C9-Cy5 and Sytox dye for 30 minutes. Supernatant of *E. coli* + C56 was supplemented with 25 µg/ml C5 conversion inhibitor OmCI to inhibit the conversion of the remaining uncleaved C5. (**A**) The percentage of bacteria with a damaged IM as determined by Sytox staining. (**B**) Deposition of C9-Cy5 on bacteria was plotted as geoMFI of the bacterial population by flow cytometry. Data represent mean +-SD of at least 3 independent experiments.

### Rapid interaction between C7 and C5b6 results in more efficient recruitment of C9

Since our data imply that assembly of MAC pores by released C5b6 does not result in similar deposition of C9 compared to by locally generated C5b6, we next wondered if rapid interaction between C7 and C5b6 at the bacterial surface enhances the recruitment of C9. Therefore, the experimental set-up of Fig. 2A was used to measure both the deposition of C9 and C6 and the ratio between these components. C9 deposition for C5b6 generated in the absence of C7 was approximately 50-fold compared to C5b6 generated in the presence of C7 (Fig. 5A). This coincided with a five-fold decrease of C6 deposition for C5b6 generated in the absence of C7 (Fig. 5B). This difference in C6 deposition was not caused by a difference in conversion of C5, as shown by similar amounts of C5a in the supernatant (**S5**). Since one C5b6 molecule is expected to recruit up to 18 C9 molecules to form a complete MAC pore (10,11), we wanted to analyze the ratio between these components based on the fluorescence measured by flow cytometry. This ratio could indicate whether or not rapid interaction between C5b6 and C7 affects the total amount of C9 molecules per C6 molecule on the surface. For MAC pores derived from C5b6 generated in the presence of C7, the C9:C6 ratio was ten-fold higher compared to C5b6 generated in the absence of C7 (Fig. 5C). Together, these data indicate that rapid interaction between C7 and C5b6 enhances the recruitment of C9 to the bacterial surface.

**Figure 5.**
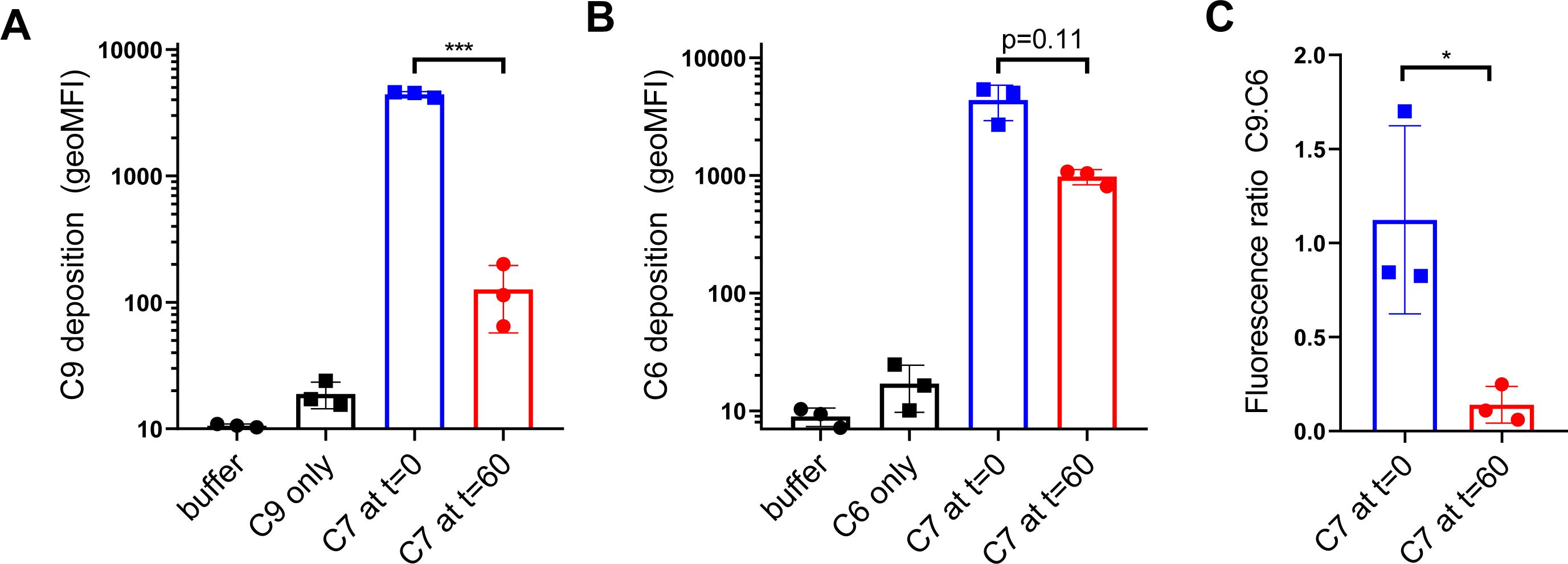
The presence of C7 during the generation of C5b6 results in more efficient recruitment of C9. Convertase-labelled *E. coli* MG1655 was treated as in Fig. 3A in which C5b6 was formed for 60 minutes in the absence (**red circles**) or presence (**blue squares**) of C7 after which C8 and C9 were added and bacteria were analyzed by flow cytometry after 30 minutes. Deposition of C9-Cy5 (**A**) and C6-Cy5 (**B**) were both measured in parallel experiments and represented as geoMFI of the bacterial population. The fluorescence ratio between C9:C6 (**C**) was calculated by dividing the geoMFI of C9 by the geoMFI of C6. Data represent individual samples with mean +-SD of at least 3 independent experiments. Statistical analysis was done using a paired one-way ANOVA with Tukey’s multiple comparisons’ test (**A-B**) and an unpaired two-tailed t-test (**C**). Significance was shown as * p ≤ 0.05, ** p ≤ 0.01, *** p ≤ 0.001 or **** p ≤ 0.0001.

### Rapid interaction between C7 and C5b6 affects how C5b-7 is anchored to the bacterial cell envelope

We next wondered if the rapid interaction between C7 and C5b6 affects how C5b-7 is anchored to the bacterial cell envelope. Recently, we showed that bactericidal MAC pores are insensitive to proteolytic cleavage by trypsin, whereas non-bactericidal MAC pores are cleaved by trypsin (16). We hypothesize that rapid interaction between C7 and C5b6 could more efficiently anchor C5b-7 to the bacterial cell envelope and render the complex insensitive to proteolytic cleavage by trypsin. Convertase-labelled bacteria were therefore labelled with C5b-7 either by locally generating C5b6 in the presence of C7 or by adding purified C5b6 (pC5b6) together with C7 and then exposed to trypsin. Next, trypsin was inhibited with soy-bean trypsin inhibitor and C8 and C9 were added to measure to measure the deposition of C9 (schematic overview Fig. 6A). When pC5b6 was used to assemble C5b-7, trypsin shaving led to a twenty-fold decrease in C9 deposition (Fig. 6B). By comparison, when C5b6 was generated locally in the presence of C7, trypsin resulted in a two-fold decrease in C9 deposition (Fig. 6B). Moreover, locally generated C5b-7 that was exposed to trypsin could still form MAC pores that damaged the bacterial IM when C8 and C9 were added (Fig. 6C), although titration of the amount of C5b-7 on the bacterial surface revealed a two-fold decrease in IM damage due to trypsin shaving.

**Figure 6.**
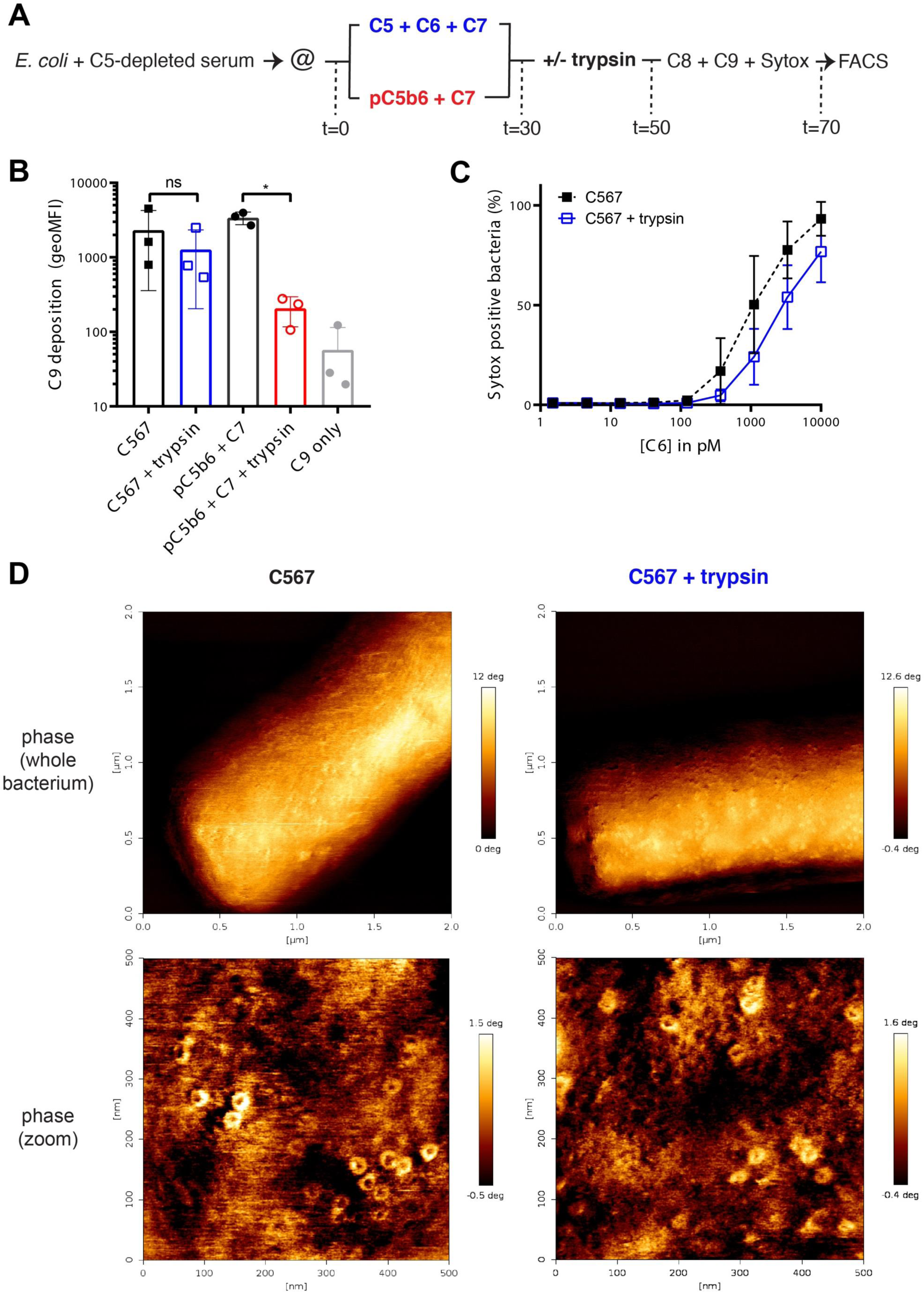
The presence of C7 during C5b6 generation affects how C5b-7 is anchored to the bacterial cell envelope. **A)** Schematic overview of trypsin shaving of *E. coli* MG1655 bacteria labelled with locally formed C5b-7 and C5b-7 derived from purified C5b6 (pC5b6). Convertase-labelled bacteria were incubated with 10 nM C5, 10 nM C6 and 10 nM C7 (**black squares**) or 10 nM pC5b6 with 10 nM C7 (**black circles**). After 30 minutes, bacteria were treated with 10 µg/ml trypsin for 20 minutes (**open blue squares** and **open red circles**). Trypsin was inhibited with 50 µg/ml soy-bean trypsin inhibitor and 10 nM C8, 100 nM C9-Cy5 and Sytox were added to complete MAC pores for 20 minutes after which bacteria were analyzed by flow cytometry. **(B)** Deposition of C9-Cy5 on bacteria treated as described in (**A**) was plotted as geoMFI of the bacterial population. (**C**) The percentage of bacteria with a damaged IM as determined by Sytox staining treated as described in (**A**). **(D)** Atomic force microscopy analysis (phase images) of *E. coli* MG1655 immobilized on Vectabond covered glass slides. Bacteria were treated as in (**A**) to label them with locally formed C5b-7 which was treated without or with trypsin before completing MAC pores by addition of C8 and C9. The concentrations of MAC components were ten-fold higher than described in (**A**). 2×2 µm^2^ scans of whole bacteria were shown on top and representative 500×500 nm^2^ zoomed images were shown on the bottom. Images are representative for a total of three bacteria per condition and at least four images per bacterium. Vertical scale bars: 12 deg for 2×2 µm^2^ scans and 2 deg for 500×500 nm^2^ scans. Data represent with mean +-SD (**B** and **C**) of at least 3 independent experiments. Statistical analysis was done using a paired one-way ANOVA with Tukey’s multiple comparisons’ test (**B**). Significance was shown as * p ≤ 0.05.

We next wanted to confirm that C9 deposition correlates with the formation of complete MAC pores. In our previous study we have shown that atomic force microscopy (AFM) can be used to detect complete MAC pores on bacteria (16). Non-bactericidal MAC pores derived from purified C5b6 could not be detected by AFM in this study, which is why we focused on analysing MAC pores derived from C5b6 generated in the presence of C7 by AFM. Therefore, bacteria were immobilized on glass slides and labelled with convertases. These convertase-labelled bacteria were incubated with C5, C6 and C7 and subsequently exposed to trypsin as in Fig. 6A. Confocal microscopy was used on non-bactericidal MAC pores derived from pC5b6 to confirm that trypsin shaving on immobilized bacteria was comparable to cells in suspension (**S6A**). Complete MAC pores were still detectable by AFM when C5b-7 was exposed to trypsin (Fig. 6D). Quantification based on all acquired images revealed that the average amount of MAC pores/µm^2^ was reduced approximately two-fold by exposing C5b-7 to trypsin (**S6B**), corresponding with the two-fold decrease in C9 deposition by flow cytometry (Fig. 6B). Altogether, these data show that rapid interaction between C7 and C5b6 is required for efficient anchoring of bactericidal C5b-7 to the bacterial cell envelope.

### Complement-resistant *E. coli* can prevent efficient anchoring of C5b-7 and insertion of MAC pores into the bacterial cell envelope

Finally, we wondered whether the above insights into MAC assembly could help understand how bacteria resist complement-dependent killing. In particular, earlier reports have shown that certain Gram-negative bacteria can resist complement-dependent killing despite the deposition of MAC pores (23–25). In line with these earlier reports, in our strain collection we also identified clinical *E. coli* strains that were complement-resistant although MAC pores were efficiently formed. Based on the molecular insights mentioned above, we wondered whether the anchoring of C5b-7 and insertion of MAC pores into the bacterial cell envelope was affected on these strains. We selected two clinical complement-resistant *E. coli* strains isolated from the urine of patients (552059.1 and 552060.1). When these clinical *E. coli* strains were incubated in C5-depleted serum we observed comparable complement activation to complement-sensitive strains (MG1655 and 547563.1), as measured by the deposition of C3b (Fig. 7A). Incubating these convertase-labelled bacteria with purified MAC components (C5-9) did not cause IM damage (Fig. 7B) despite comparable deposition of C9 to complement-sensitive strains (Fig. 7C). Next, MAC pores on these bacteria were shaved with trypsin after MAC pores were completed to measure if MAC pores were efficiently inserted into the bacterial cell envelope. Trypsin shaving of MAC pores in both complement-resistant strains resulted a 90% decrease of C9 deposited on the surface compared to a no trypsin control, whereas on complement-sensitive strains this decrease was only 20 or 40% for MG1655 and 547563.1, respectively (Fig. 7D). Finally, we assessed if anchoring of C5b-7 was also affected on these complement-resistant strains. Bacteria were therefore incubated with C8-depleted serum to label them with C5b-7, after which bacteria were exposed to trypsin. Next, C8 and C9-Cy5 were added so we could measure C9 deposition. On both complement-resistant strains, trypsin shaving of C5b-7 resulted in and 80% decrease in C9 deposition compared to the buffer control, whereas on complement-sensitive strains decrease in C9 deposition was 40 or 30% compared to the buffer control for MG1655 and 547563.1, respectively (Fig. 7E). Altogether, these data indicate that complement-resistant *E. coli* can prevent complement-dependent killing by MAC pores by preventing efficient anchoring of C5b-7 and insertion of MAC pores into the bacterial cell envelope.

**Figure 7.**
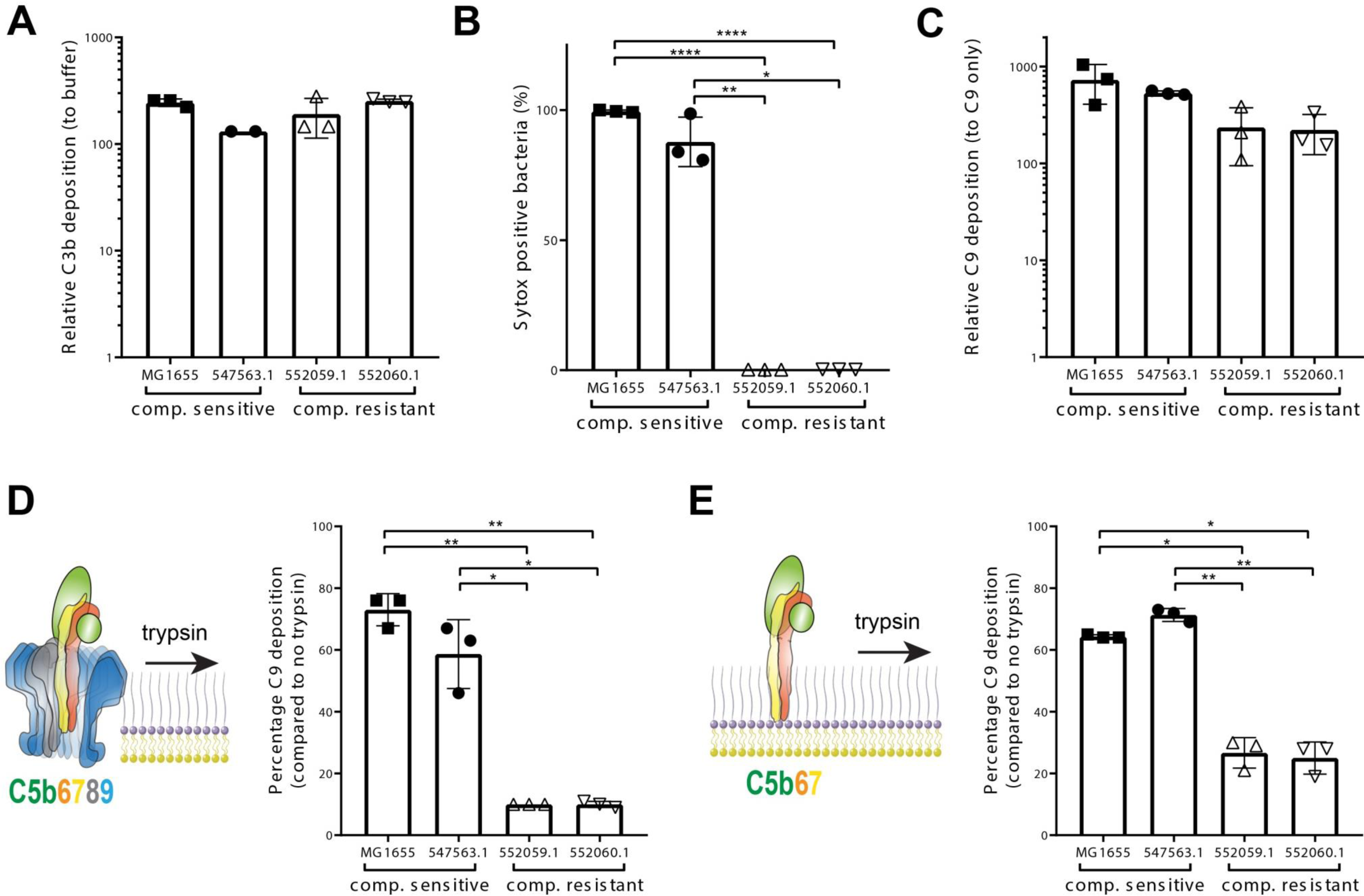
Complement-resistant *E. coli* can prevent efficient anchoring of C5b-7 and insertion of MAC pores into the bacterial cell envelope. (**A**) C3b deposition was measured for complement-sensitive (MG1655, filled squares, and 547563.1, filled circles) and complement-resistant (552059.1, open upward triangles, and 552060.1, open downward triangles) *E. coli* strains treated with 10% C5-depleted serum. C3b deposition was plotted as relative fluorescence by dividing the geoMFI of the bacterial population by the geoMFI of a control sample without antibody. Convertase-labelled bacteria were incubated with purified MAC components (C5-9) with Sytox blue to determine the percentage of Sytox positive bacteria (**B**) and the deposition of C9-Cy5 (**C**) by flow cytometry. C9-Cy5 deposition was plotted as relative fluorescence by dividing the geoMFI of the bacterial population by the geoMFI of bacteria incubated with C9-Cy5 only. Trypsin shaving experiments were performed on C5b-7 labelled bacteria by incubation in 10% C8-depleted serum. (**D**) C8 and C9-Cy5 were first added to C5b-7 labelled bacteria to assemble MAC pores, after which bacteria were treated either with buffer or 10 µg/ml trypsin. C9-Cy5 deposition was measured by flow cytometry. (**E**) C5b-7 labelled bacteria were first treated with 10 µg/ml trypsin after which 50 µg/ml soy-bean trypsin inhibitor was added together with C8 and C9-Cy5 to measure C9-Cy5 deposition by flow cytometry. For both (D) and (E), C9 deposition after trypsin treatment was plotted as the percentage of the C9 deposition compared to bacteria treated with buffer. This was done by dividing the geoMFI (Cy5) of the bacteria treated with trypsin by the geoMFI (Cy5) of bacteria treated with buffer. Data represent mean +-SD of at least 3 independent experiments (547563.1 in A was absent during one experiment, resulting in only two measurements for this strain). Statistical analysis was done using an ordinary one-way ANOVA (**A**) or paired one-way ANOVA (**B, C, D** and **E**) with Tukey’s multiple comparisons’ test. Significance was shown as * p ≤ 0.05, ** p ≤ 0.01 or **** p ≤ 0.0001.

## Discussion

A better understanding of how the immune system kills bacteria and how pathogens can counteract this may hold important clues for future control of bacterial infections. By studying how MAC pores assemble on Gram-negative bacteria, we here provide new molecular insights into the step-wise assembly of MAC pores on bacteria. Although an immediate interaction between C5b6 and C7 is not required for MAC-mediated lysis of erythrocytes (7,12,15,19,20), our data indicate that rapid interaction of C7 with convertase-generated C5b6 is essential to form bactericidal MAC pores. This is, to the best of our knowledge, the first study showing that a time-sensitive interaction between C5b6 and C7 is crucial to form functional MAC pores on bacteria. Studies on erythrocytes and liposomes have suggested that C5b6 could release from the convertase and, upon interaction with C7 in the fluid-phase, again anchor to membranes and form functional MAC pores (8,21,26–28). However, our data indicate that for the assembly of bactericidal MAC pores C5b-7 requires direct anchoring to the bacterial cell envelope to retain its bactericidal potential (Fig. 8A). This direct anchoring of C5b-7 prevents release of C5b6 from the bacterial surface, and our data indicate that released C5b6 fails to form bactericidal MAC pores despite being capable of binding to bacteria and recruiting downstream MAC components (Fig. 8B-I). This supports our earlier results showing that commercially available C5b6 (formed on zymosan surfaces) also lacks bactericidal capacity (16). Thus, in addition to the fact that local assembly of C5b6 by a C5 convertase is essential for bacterial killing (16), we now show that even locally formed C5b6 can lose its bactericidal capacity if it does not rapidly interact with C7 (Fig. 8B-II). Because this delayed interaction with C7 also correlated with an increased sensitivity of C5b-7 to trypsin, we hypothesize that the rapid interaction between C5b6 and C7 is crucial for efficient anchoring of C5b-7 to the bacterial cell envelope (Fig. 8B-III).

**Figure 8.**
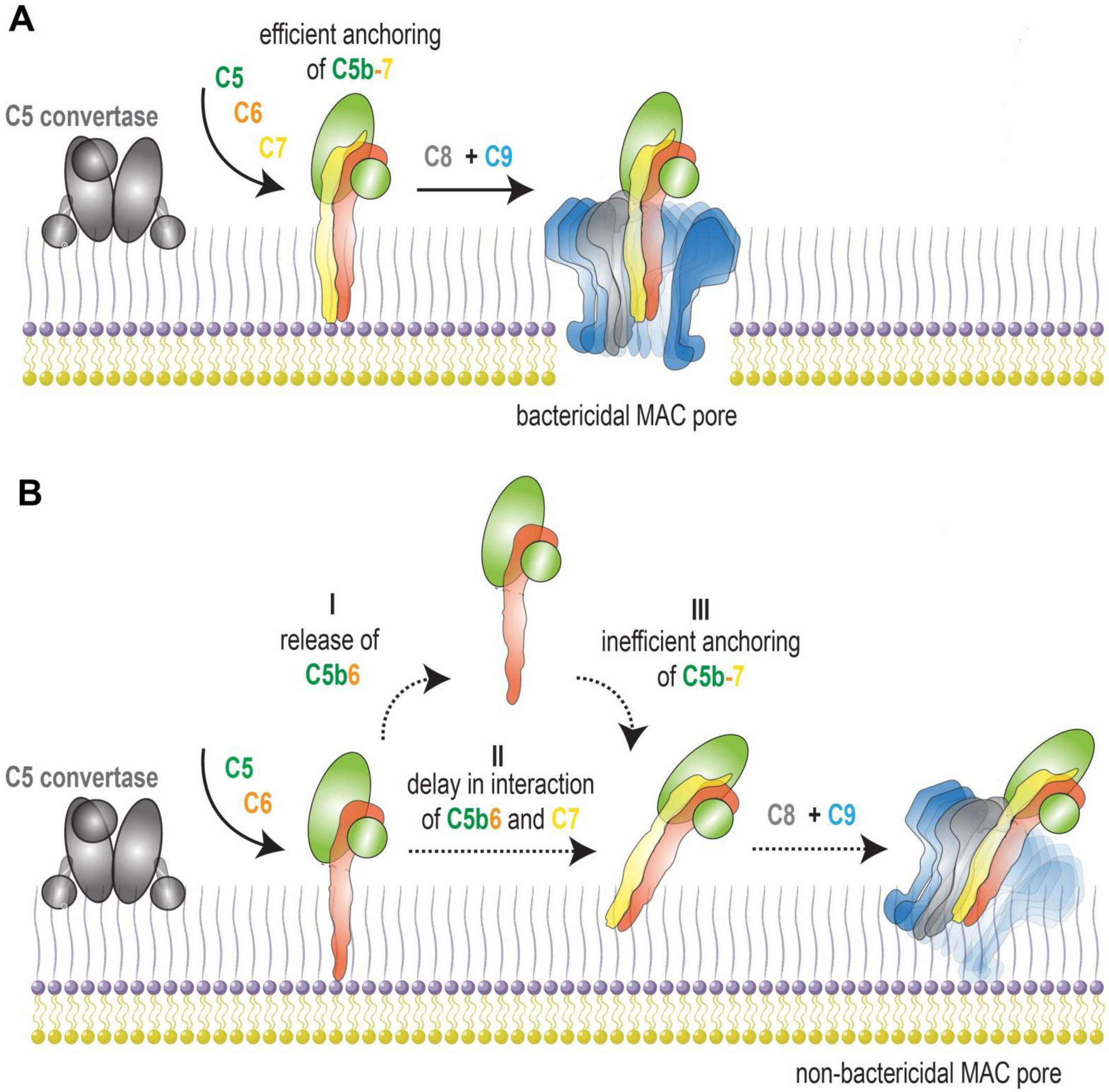
The relevance of direct assembly and anchoring of C5b-7 to the bacterial surface for bacterial killing by MAC pores. (A) Illustration depicting the relevance of direct assembly and anchoring of C5b-7 to form a bactericidal MAC pore. A surface-bound C5 convertase converts C5 into C5b in the presence of C6 and C7 to directly assembly C5b-7 and anchor C5b-7 to the bacterial surface. Subsequent recruitment of C8 and multiple copies of C9 assembles a stably inserted MAC pore that kills bacteria. (B) Formation of C5b6 in the absence of C7 can lead to the release of C5b6 (I) and/or a delay in the interaction between C5b6 and C7 (II). Both result in inefficient anchoring of C5b-7 to the bacterial surface (III). Inefficiently anchored C5b-7 can recruit C8 and C9, but these MAC pores fail to kill bacteria.

Why direct anchoring of C5b-7 is crucial for bacterial killing by MAC pores remains unclear. We hypothesize that rapid interaction of C5b6 with C7 might facilitate a specific orientation of C5b-7 that allows the metastable lipophilic domain of C7 (7) to efficiently anchor to the bacterial OM. This could be relevant for insertion of MAC pores into the OM of the bacterial cell envelope which is densely packed with outer membrane proteins (OMPs) and lipopolysaccharides (LPS). Potentially these membrane structures could also hamper efficient anchoring of C5b-7 to the OM if it is not anchored directly after the generation of C5b6 by a surface-bound convertase. Especially in the situation where C5b6 is first released from the surface, C5b6 could bind back further away from the membrane to the O-antigen of LPS or to a more proteinaceous part of the membrane or and thereby fail to efficiently anchor of C5b-7 to lipid-rich patches of the bacterial OM where the MAC pore can efficiently insert.

Our data also suggest that direct anchoring of C5b-7 to the bacterial cell envelope enhances the efficiency by which complete MAC pores are formed. Direct assembly of C5b-7 increased the ratio of C9:C6 on the bacterial surface about ten-fold, suggesting that for bactericidal MAC pores, more C9 molecules were recruited per C5b-8 complex. We do show here that the amount of MAC pores as detected by AFM correlated with amount of deposited C9 as detected by fluorescence in the case of bactericidal MAC pores. However, no such correlation was observed for non-bactericidal MAC pores in previous work since no pores could be detected by AFM (16). Given the difference in C9:C6 ratio for bactericidal versus non-bactericidal MAC pores, this could suggest that non-bactericidal MAC pores contain less C9 molecules per C5b-8 complex, which could explain the absence of detection by AFM. In experiments on model lipid membranes, MAC assembly seems rapidly completed once the first C9 is bound to C5b-8 complexes, leading to pores that are most homogeneous in size (21). However, there have been previous reports that showed heterogeneity in the amount of C9 molecules that are incorporated per C5b-8 complex (28–31). Moreover, others have also compared C9 polymerization or the ratio of C9 to C5b-8 complexes for bactericidal and non-bactericidal MAC pores, but so far conflicting results have made it hard to draw conclusions on how polymerization of C9 affects bacterial killing (3,23,24,32).

Finally, by studying MAC assembly on clinical, complement-resistant bacteria, we revealed that Gram-negative bacteria have evolved mechanisms to resist killing by MAC pores, despite being efficiently recognized by complement and labelled with functional C5 convertases. We identified complement-resistant *E. coli* strains that blocked MAC-mediated killing by preventing efficient anchoring of C5b-7 to the bacterial cell envelope and subsequent insertion of MAC pores. Although several mechanisms of bacterial complement evasion have been described (33–36), these are mostly not MAC-specific because they block initial complement activation steps such as recognition or the deposition of functional convertases (35,37,38). Preventing efficient anchoring of C5b-7 to thus affect insertion of MAC pores into the bacterial cell envelope has not, to our knowledge, been reported before on *E. coli*. Others have previously reported inefficient anchoring of C5b-7 on a complement-resistant *Salmonella minnesota* strain (23,39). How Gram-negative bacteria can prevent C5b-7 anchoring remains unclear, but modifying structures covering the target membrane could hamper this crucial step. Modifications to capsular polysaccharides and LPS in gonococci have previously been linked to complement-resistance (40). However, these modifications could also affect initial recognition and complement activation, which was not observed in the *E. coli* strains used in this study. It is interesting to note we and others have also shown abundant deposition of MAC components on several Gram-positive bacteria (41,42), while these are intrinsically resistant against killing by the MAC. For Gram-positives it is likely the thick peptidoglycan layer that specifically prevents insertion of MAC pores in the cytosolic membrane.

Altogether this study provides a better molecular understanding of how MAC pores have to be assembled to elicit bacterial killing. Understanding how bacteria are killed by MAC pores and how bacteria can counteract this complement-mediated killing is important to underpin how bacteria escape the immune system and cause infections.

## Acknowledgements

This work was funded by: an ERC Starting grant (639209-ComBact, to S.H.M.R), the Utrecht University Molecular immunology HUB (eSTIMATE) and the UK EPSRC and MRC (EP/N509577/1 and MR/R000328/1).

## Materials & Methods

### Serum and complement proteins

Normal human serum (NHS) was obtained from healthy volunteers as previously described (43). Serum depleted of complement components C5, C6 or C8 was obtained from Complement Technology. Complement components purified C5b6 (pC5b6) and C8 were also obtained from Complement Technology. His-tagged complement components C5, C6, C7, C9 and were expressed in HEK293E cells at U-Protein Express and purified on HisTrap Excel columns. OmCI was produced in HEK293E cells at U-Protein Express as well and purified as described before (44). To produce fluorescently labelled complement components C6 and C9, proteins were modified with a C-terminal LPETG-His tag and expressed and purified as other His-tagged complement components. 50 µM of LPETG-His tagged protein was incubated with 1 mM GGG-azide (Genscript) and 25 µM His-tagged sortase-A7+ (recombinantly expressed in *E. coli*) for 2h at 4 °C in Tris/NaCl buffer (50 mM Tris/300 mM NaCl and the tagged protein was purified over a HisTrap FF column (GE Healthcare). GGG-azide labelled proteins were washed with Tris/NaCl buffer and concentrated to 25 µM on a 30 kDa Amicon Tube (Merck Millipore) and afterwards labelled with 100 µM DBCO-Cy5 (Sigma Aldrich) via copper-free click chemistry for 3h at 4 °C. Finally, Cy5-labelled proteins were purified by size-exclusion chromatography (SEC) on a Superdex 200 Increase (GE Healthcare) column with PBS and concentrated on a 30 kDa Amicon Tube. Labelling of the proteins was verified by measuring OD633 during SEC on the Akta Explorer (GE Healthcare). The final concentration was determined by OD280 measurement on the Nanodrop and verified by SDS-PAGE.

### Bacterial strains

Unless otherwise specified, a common laboratory *E. coli* strain MG1655 was used in our experiments. Other common laboratory *E. coli* strains that were used were BW25113 and MC1061. *E. coli* strain 547563.1, 552059.1 and 552060.1 were clinical isolates all isolated from urine and obtained from the clinical Medical Microbiology department at the University Medical Center Utrecht.

### Bacterial growth

For all experiments, bacteria were plated on Lysogeny Broth (LB) agar plates. Single colonies were picked and grown overnight at 37 °C in LB medium. The next day, subcultures were grown by diluting at least 1/30 and these were grown to mid-log phase (OD600 between 0.4 – 0.6). Once grown to mid-log phase, bacteria were washed by centrifugation three times (11000 rcf for 2 minutes) and resuspended to OD 1.0 (~1 × 10^9^ bacteria/ml) in RPMI (Gibco) + 0.05% human serum albumin (HSA, Sanquin).

### Complement-dependent bactericidal & complement deposition assays

In all experiments, bacteria were labelled with convertases as described previously in Heesterbeek *et al*. (16) unless stated differently. In short, bacteria at OD600 = 0.1 (~1 × 10^8^ bacteria/ml) were incubated with 10% C5-depleted serum at 37 °C, washed three times and resuspended RPMI-HSA. C3b deposition was measured by incubating bacteria at OD600 = 0.05 (~5 × 10^7^ bacteria/ml) with 3 µg/ml mouse-anti C3b labelled with Alexa Fluor A488 (previously described in Heesterbeek et al (16)) for 30 minutes at 4 °C. Convertase-labelled bacteria were incubated at OD600 = 0.05 (~5 × 10^7^ bacteria/ml) with complement components at 37 °C. Unless stated differently, the concentration of complement components C5, C6, C7, C8 and C9 was 10/10/10/10/20 nM respectively. These concentrations of complement components reflect the concentration of these components in 1 – 3% NHS (100% NHS has ~ 375 nM C5, 550 nM C6, 600 nM C7, 350 nM C8 and 900 nM C9). When pC5b6 was used in an experiment, the concentration of pC5b6 was 10 nM and of C9 100 nM. In experiments where OmCI was used, 25 µg/ml (~ 1 µM) was added. For experiments where complement components were added at different points in time, the total incubation time is described in the figure legends. For experiments in which bacteria were first labelled with C5b-7, labelling of C5b-7 was performed for 30 minutes at 37 °C and were subsequently washed three times. If bacteria were labelled with C5b-7 via purified components, bacteria were first mixed with C5, C6 or pC5b6 before adding C7 to prevent fluid-phase aggregation of C5b-7 complexes. When bacteria were immediately labelled with both convertases and C5b-7 in one step, bacteria were incubated with 10% C8-depleted serum at OD600 = 1.0 (~1 × 10^9^ bacteria/ml) for 30 minutes at 37 °C. C5b-7 labelled bacteria were subsequently incubated with C8 and C9 for 30 minutes at 37 °C. In all experiments, 2.5 µM of Sytox Blue Dead Cell stain (Thermofisher) was added to the final incubation step of the experiment. Samples were diluted 10-20 times and subsequently analyzed in the BD FACSVerse flow cytometer for forward scatter (FSC), side scatter (SSC), Sytox and Cy5 intensity. In all these experiments, RPMI + 0.05% HSA was used as buffer and all washing steps were done by centrifugation at 11000 rcf for 2 minutes.

### Collecting supernatant containing released C5b6

For assays in which bacterial supernatants were harvested, bacteria were labelled with convertases as described previously with C5-depleted serum with RPMI + 0.05% HSA as buffer. In these assays, the bacterial concentration and complement component concentration was increased ten-fold to maximize the yield of released C5b6. Therefore, bacteria at OD600 = 0.5 (~5 × 10^8^ bacteria/ml) and C5, C6, C7, C8 and C9 at 100/100/100/100/1000 nM were incubated for 1h at 37 °C. Bacteria were next spun down to collect the supernatant, which was supplemented with 25 µg/ml OmCI (~ 1 µM) to specifically block uncleaved C5 in further assays.

### C5b Western blots

Bacterial supernatants were collected as described above and the cell pellets were also collected. For reducing SDS-PAGE, samples were diluted 1:1 in 2x SDS sample buffer (0.1M Tris (pH 6.8), 39% glycerol, 0.6% SDS and bromophenol blue) supplemented with 50 mg/ml dithiothreitol (DTT) and placed at 95 °C for 5 minutes. Samples were run on a 4-12% Bis-Tris gradient gel (Invitrogen) for 60 minutes at 200V. Proteins were subsequently transferred with the Trans-Blot Turbo Transfer system (Bio-Rad) to 0.2 µM PVDF membranes (Bio-Rad). Initially, samples were blocked with PBS supplemented with 0.1% Tween-20 (PBS-T) and 4% dried skim milk (ELK, Campina) for 60 minutes at 37 °C. Primary staining was performed with a 1:500 dilution (~ 80 µg/ml) of polyclonal goat-anti human C5 (Complement Technology) in PBS-T supplemented with 1% ELK for 60 minutes at 37 °C. Secondary staining was performed with a 1:10.000 of HRP-conjugated pooled donkey antisera against goat IgG (H+L) (Southern Biotech) in PBS-T supplemented with 1% ELK for 60 minutes at 37 °C. In between all steps and after the final staining, membranes were washed three times with PBS-T. Finally, membranes were developed with Pierce ECL Western Blotting Substrate (Thermo Scientific) for 1 minute at RT and imaged on the LAS4000 Imagequant (GE Healthcare).

### C5b6 ELISA

Bacterial supernatants were collected as previously described. Nunc Maxisorp ELISA plates were coated overnight at 4 °C with 50 µl/well of 3 µg/ml monoclonal mouse IgG1 anti human C6 (Quidel) in PBS. Blocking was performed with PBS-T supplemented with 4% bovine serum albumin (BSA, Sigma) for 60 minutes at 37 °C. Dilutions of supernatant and standard of 1 nM pC5b6, C5 or C6 were prepared in PBS-T supplemented with 1% BSA and were bound for 60 minutes at 37 °C. Primary staining was performed with a 1:500 dilution (~ 80 µg/ml) of polyclonal goat-anti human C5 (Complement Technology) in PBS-T supplemented with 1% BSA for 60 minutes at 37 °C. Secondary staining was performed with a 1:5.000 of HRP-conjugated pooled donkey antisera against goat IgG (H+L) (Southern Biotech) in PBS-T supplemented with 1% BSA for 60 minutes at 37 °C. In between all steps, wells were washed three times with PBS-T. Finally, fresh tetramethylbenzidine (TMB) was added for development, the reaction was stopped with 4N sulfuric acid and the OD450 was measured.

### Haemolytic assays

Rabbit erythrocytes were collected from rabbit blood obtained from the Veterinary Department and stored 1:1 in Alsever (0.4% NaCl, 2% D-glucose, 0.8% sodium citrate, 0.055% citric acid) for a maximum of two weeks. Rabbit erythrocytes were isolated by spinning down the blood at 1.000 rcf for 5 minutes and washed three times with PBS after which the supernatant was discarded. The packed erythrocyte pellet was resuspended in Veronal buffered saline (2 mM Veronal, 145 mM NaCl, pH = 7.4) supplemented with 0.1% BSA and 2.5 mM MgCl2 (VBS+), which was the buffer that was maintained during the experiments. For experiments with purified C5b6 or released C5b6 in supernatants, both rabbit and human erythrocytes (4%, 1×10^8^/ml) were incubated with a titration of C5b6 together with 10 nM C7, 10 nM C8 and 100 nM C9 for 30 minutes at 37 °C. The mixture of C7, C8 and C9 was always added last to avoid fluid-phase formation of MAC complexes. Haemoglobulin release was measured by spinning down the plate (1250 rcf for 5 minutes) and subsequently diluting the supernatant 1:3 in MilliQ (MQ) and measuring the absorbance at OD405 nm. MQ was used as positive control for maximum lysis of erythrocytes. The percentage of erythrocyte lysis was subsequently calculated by setting buffer control at 0% lysis and MQ control at 100% lysis.

### Trypsin shaving

Bacteria were labelled with C5b-7 as described previously and subsequently washed three times by centrifugation (11000 rcf for 2 minutes). Bacteria were resuspended into RPMI + 0.05% HSA, which was used as buffer for all other incubations. For experiments where C5b-7 was shaved with trypsin, C5b-7 labelled bacteria at OD600 = 0.05 (5×10^7^/ml) were incubated with 10 µg/ml bovine trypsin (Sigma) for 20 minutes at 37 °C. Next, 25 µg/ml soy bean trypsin inhibitor (sbti, Sigma) was added along with 10 nM C8 and 20 nM C9 for 20 minutes at 37 °C. For experiments where C5b-9 was trypsinized, C5b-7 labelled bacteria were first incubated with 10 nM C8 and 20 nM C9 for 20 minutes at 37 °C and then exposed to 10 µg/ml trypsin for 20 minutes at 37 °C. In all experiments, 2.5 µM of Sytox Blue Dead Cell stain (Thermofisher) was added to the final incubation step of the experiment. Samples were diluted 10-20 times and subsequently analyzed in the BD FACSVerse flow cytometer for FSC, SSC, Sytox, GFP and Cy5 intensity.

### Atomic force microscopy

Bacteria were grown as previously described and resuspended in HEPES buffered saline (HBS, 20 mM HEPES/120 mM NaCl at pH 7.4) to OD600 = 1.0 (1×10^9^ bacteria/ml). 100 µl of bacteria were immobilized on cleaned glass slides (Corning/Sigma Aldrich) covered with Vectabond^®^ (Vector Laboratories, USA) as described previously for 30 minutes at RT (45). During all steps, care was taken in preventing the droplet from drying out. Immobilized bacteria were labelled with convertases by incubation with 10% C5-depleted serum in HBS supplemented with 0.1% BSA and 2.5 mM MgCl_2_ (HBS+) for 20 minutes at 37 °C. Immobilized bacteria were then washed with HBS+ and treated with 100 nM C5, 100 nM C6 and 100 nM C7 in HBS+ for 30 minutes at 37 °C. Next, immobilized bacteria were washed and treated with 10 µg/ml trypsin in HBS+ for 30 minutes at 37 °C. This was followed by a wash and treatment with 100 nM C8 and 1000 nM C9 for 30 minutes at 37 °C. Finally, bacteria were washed with HBS and imaged by AFM.

Intermittent-contact mode atomic force microscopy was performed on a Nanowizard III AFM with an UltraSpeed head (Bruker AXS, CA, USA) and a FastScanD (Bruker AXS, CA, USA) cantilever with 0.25 N/m spring constant and 120 kHz resonance frequency. Drive frequencies were between 95 and 125 kHz and amplitudes were between 8.5-20.5 nm, equivalent to a 10-50% drop from the free amplitude 5-10 µm from the sample surface. AFM was performed in HBS within 4 hours of sample preparation. Images are 512×512 pixels. Whole bacteria scans were 2×2 µm^2^; 500×500 nm^2^ scans were then performed over the bacterial surface. Large scans were performed at 1-3 Hz and smaller scans at 10-15 Hz. Data was analysed in JPK data processing software (Bruker AXS, CA, USA). 2×2 µm^2^ scans were processed by applying a first-order plane fit. 500×500 nm^2^ scans had a line-by-line 2^nd^ or 3^rd^ order flattening and a Gaussian filter with σ = 1 pixel, to remove high-frequency noise, applied.

### Data analysis and statistical testing

Flow cytometry data was analysed in FlowJo V.10. MG1655 bacteria were gated on FSC, SSC and being GFP-positive. All other bacteria were gated based on FSC and SSC only. Sytox positive cells were gated such that the buffer only control had <1 % positive cells. Unless stated otherwise, graphs are comprised of at least three biological replicates. Statistical analyses were performed in GraphPad Prism 8 and are further specified in the figure legends.

