## Supplemental information for "Bacterial killing by complement requires direct anchoring of Membrane Attack Complex precursor C5b-7"

**S1 – FACS gating strategy and histograms**

Gating strategy of bacteria analysed by flow cytometry. Bacteria were gated based on forward scatter and side scatter. Sytox positive cells were gated such that the buffer control (top right) had <1% positive cells. Representative plots showing convertase-labelled MG1655 incubated with buffer only, C5-9 at t=0, C7 at t=0 and C7 at t=60 samples described in **Fig. 2A**.

**S2 – Validation specificity of C5b6 ELISA**

Specificity of the C5b6 ELISA is shown here. A titration of C5, C6, C5 + C6, purified C5b6 (pC5b6) or supernatant of convertase-labelled *E. coli* MG1655 incubated with C5 + C6 was added to ELISA plates coated with monoclonal anti-human C6 and next detected with polyclonal anti-C5. Data represent mean +- SD of at least 3 independent experiments.

**S3 – Functionality C6-Cy5**

*E. coli* MG1655 bacteria were added to 1% C6-depleted serum supplemented with a titration of C6 isolated from plasma (CT = Complement Technology), recombinantly expressed C6-LPETG-His (LPETG) and sortagged C6-LPETGGGG-Cy5 (Cy5). (**A**) The percentage of bacteria with a damaged inner membrane as determined by Sytox staining. **(B)** Deposition of C6-Cy5 on bacteria was plotted as geoMFI of the bacterial population. Data represent mean +- SD of at least 3 independent experiments.

**S4 – Rabbit erythrocyte lysis of released C5b6 in bacterial supernatant**

Rabbit erythrocytes were incubated with a titration of purified C5b6 (pC5b6) or supernatant of convertase-labelled *E. coli* MG1655 incubated with C5 + C6 in the presence of 10 nM C7, 10 nM C8 and 100 nM C9. The percentage of erythrocytes that were lysed was substracting background OD405 of erythrocytes in buffer (0% lysis) from each value and dividing this by the OD405 value of erythrocytes in MilliQ (100% lysis). Data represent mean +- SD of at least 3 independent experiments.

**S5 – C5a generation in the presence or absence of C7**

C5a was measured in the supernatant of convertase-labelled *E. coli* MG1655 incubated with 100 nM C5, 100 nM C6 in the absence (**red circles**) or presence (**blue squares**) of 100 nM C7 by a calcium flux-based reporter assay (46). Supernatant was diluted 1/30, 1/100 and 1/300 times. A titration of purified C5a (filled triangles) was taken as standard. Data represent mean +- SD of at least 3 independent experiments.

**S6 – Validation trypsin shaving on glass slides & quantification MAC pores by atomic force microscopy**

(A) GFP-induced *E. coli* MG1655 were immobilized on Cell tak (BD Diagnostics, USA) covered glass slides and next labelled with convertases with 10% C5-depleted serum. Next, bacteria were incubated with 100 nM pC5b6 , 100 nM C7, 100 nM C8 and 1000 nM C9 and subsequently treated with buffer or 10 µg/ml trypsin. Samples were imaged using a Leica SP5 confocal microscope with a HCX PL APO CS 63x/1.40-0.60 OIL objective (Leica Microsystems, the Netherlands). (**B**) Quantification of MAC pores on atomic force microscopy images (phase images) of *E. coli* MG1655 immobilized on Vectabond covered glass slides and treated as in **Fig 6D**. The number of MACs per 500x500 $nm^2$ scan was counted by hand and used to calculate the number of MACs per $\mu m^2$ for each analyzed bacterium. Three bacteria were examined in each condition with at least four smaller scans per cell.

**S1**

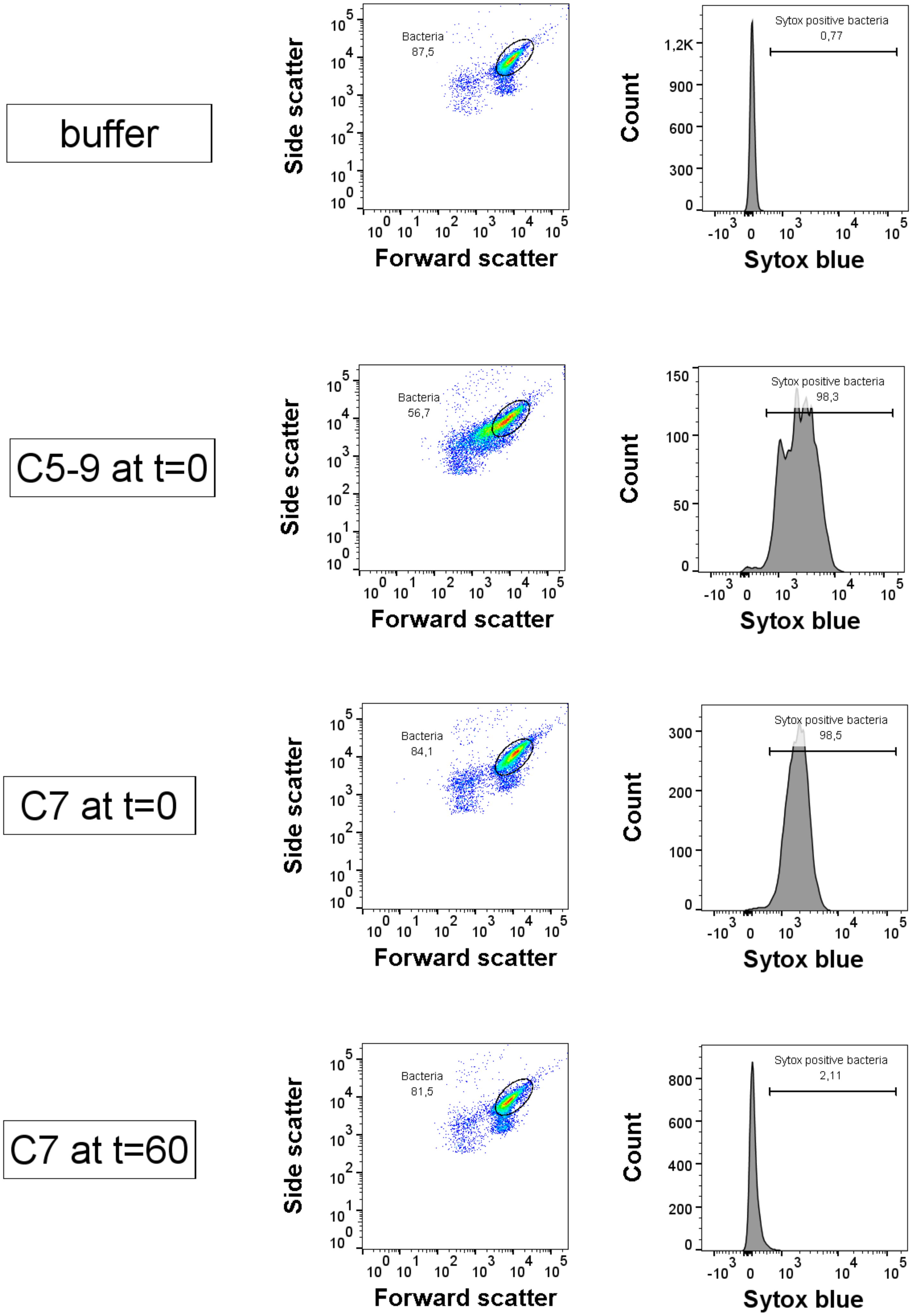
buffer
Bacteria
87,5
Sytox positive bacteria
0,77
C5-9 at t=0
Bacteria
56,7
Sytox positive bacteria
98,3
C7 at t=0
Bacteria
84,1
Sytox positive bacteria
98,5
C7 at t=60
Bacteria
81,5
Sytox positive bacteria
2,11
Side scatter
Forward scatter
Count
Sytox blue

**S2**

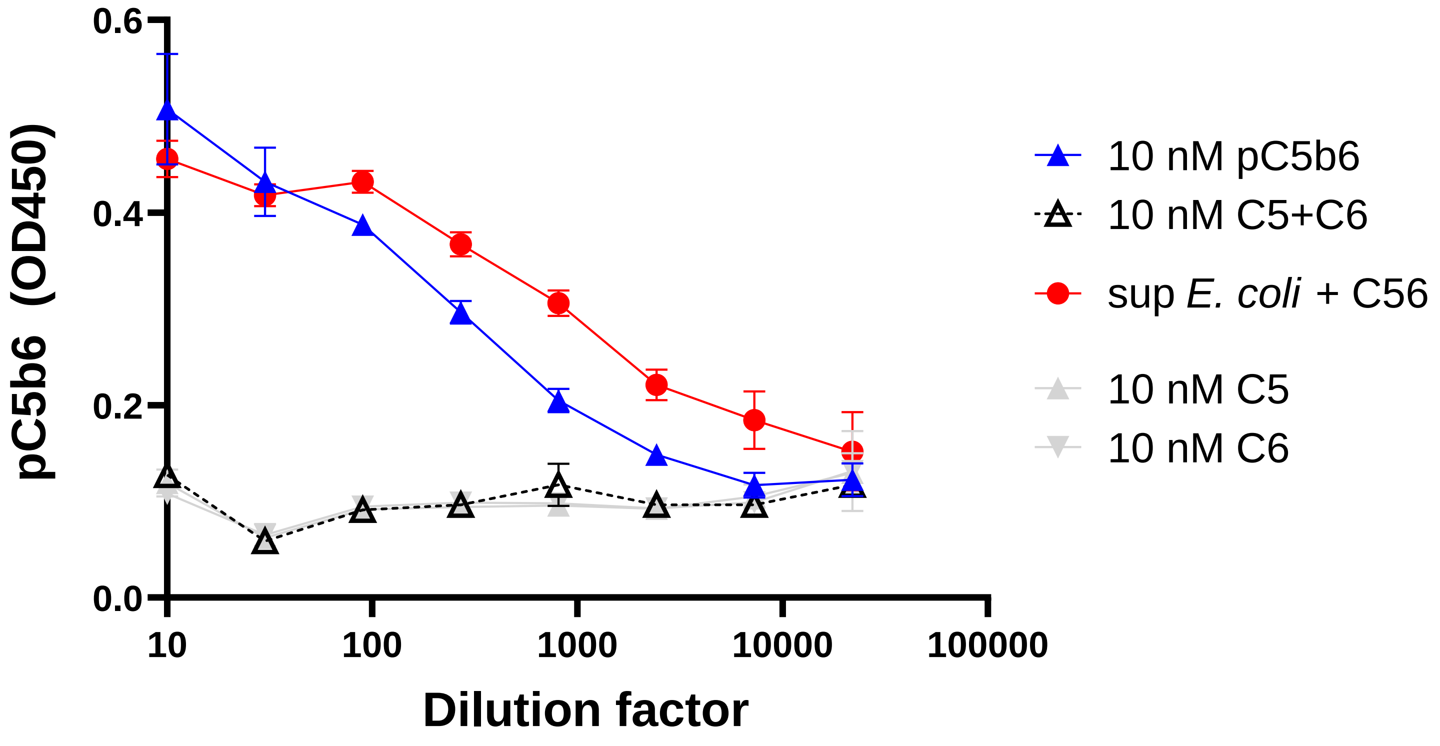

pC5b6 (OD450)
0.6
0.4
0.2
0.0
10
100
1000
10000
100000
Dilution factor
10 nM pC5b6
10 nM C5+C6
sup *E. coli* + C56
10 nM C5
10 nM C6

## S3

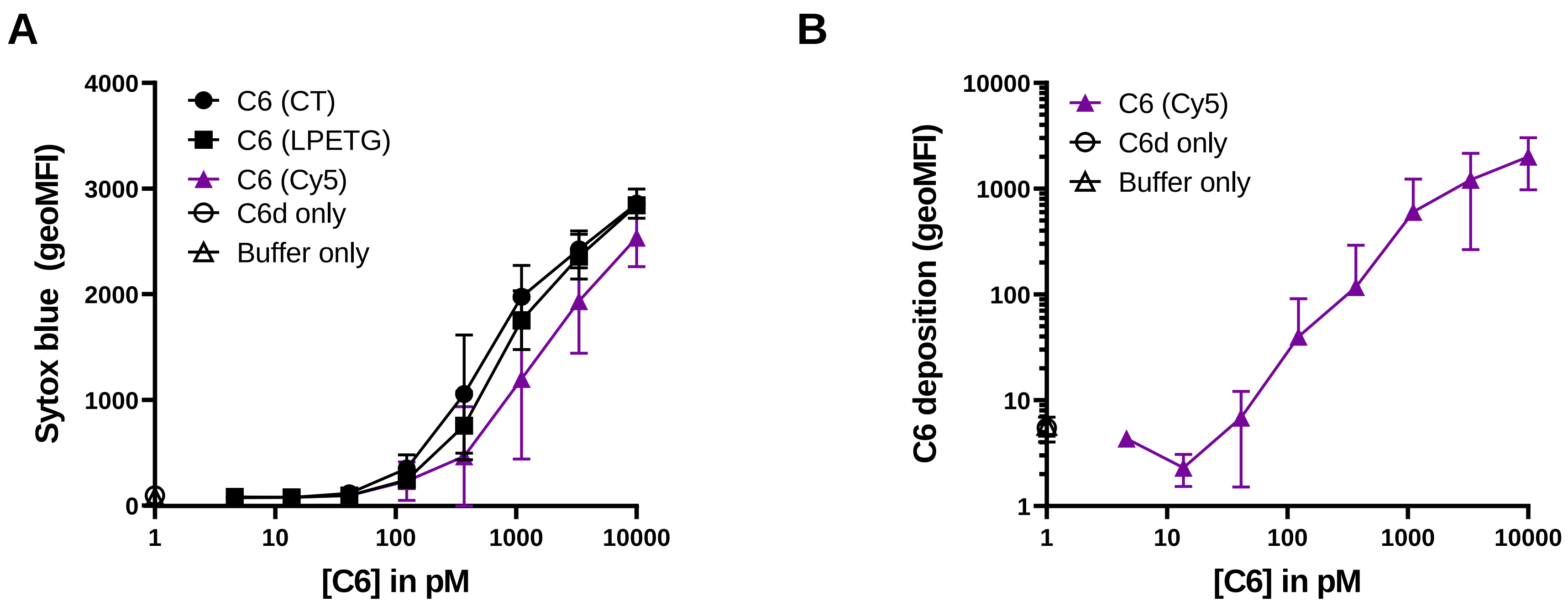

A
B
Sytox blue (geoMFI)
4000
3000
2000
1000
0
1
10
100
1000
10000
[C6] in pM
C6 (CT)
C6 (LPETG)
C6 (Cy5)
C6d only
Buffer only
C6 deposition (geoMFI)
10000
1000
100
10
1
C6 (Cy5)
C6d only
Buffer only

**S4**

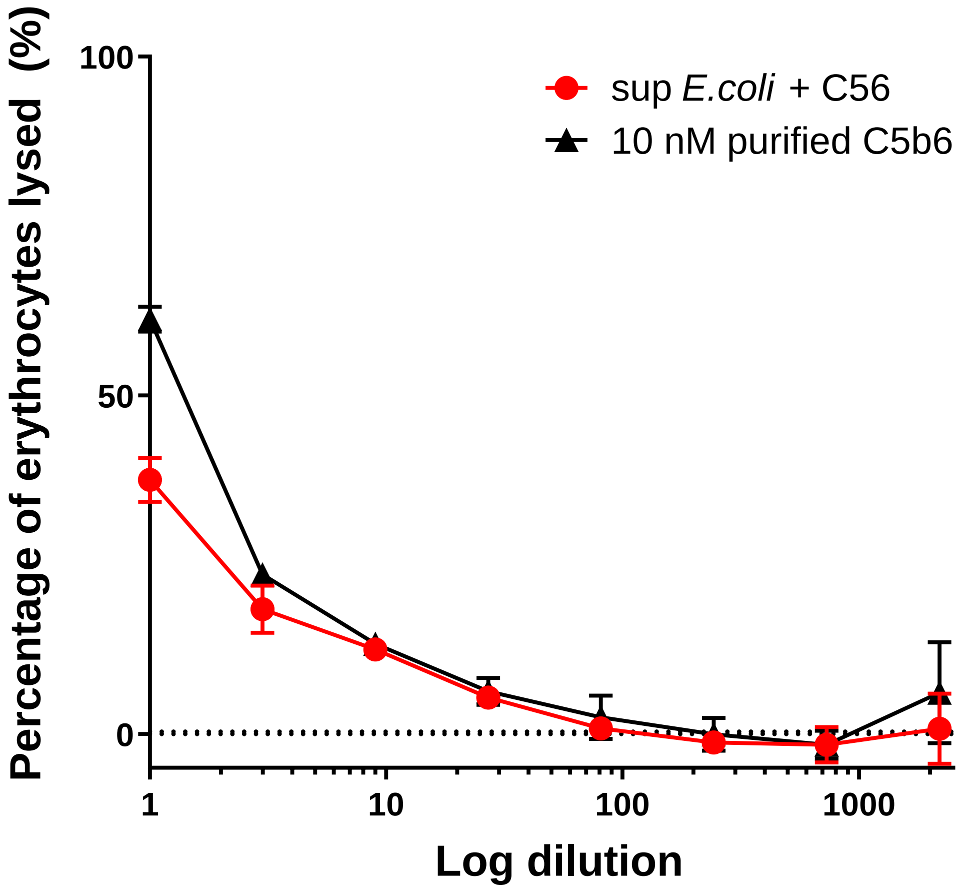

sup *E.coli* + C56
10 nM purified C5b6
Percentage of erythrocytes lysed (%)
100
50
0
1
10
100
1000
Log dilution

**S5**

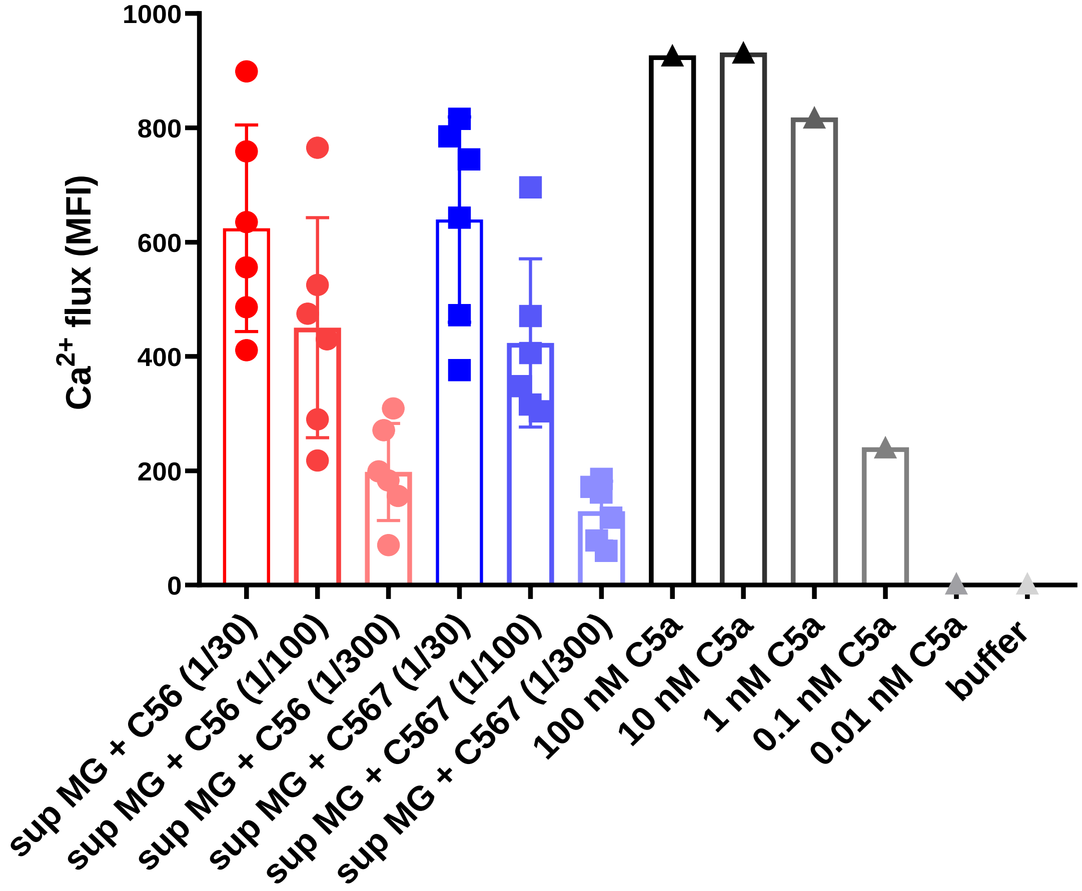

Ca2+ flux (MFI)
1000
800
600
400
200
0
sup MG + C56 (1/30)
sup MG + C56 (1/100)
sup MG + C56 (1/300)
sup MG + C567 (1/30)
sup MG + C567 (1/100)
sup MG + C567 (1/300)
100 nM C5a
10 nM C5a
1 nM C5a
0.1 nM C5a
0.01 nM C5a
buffer

S6

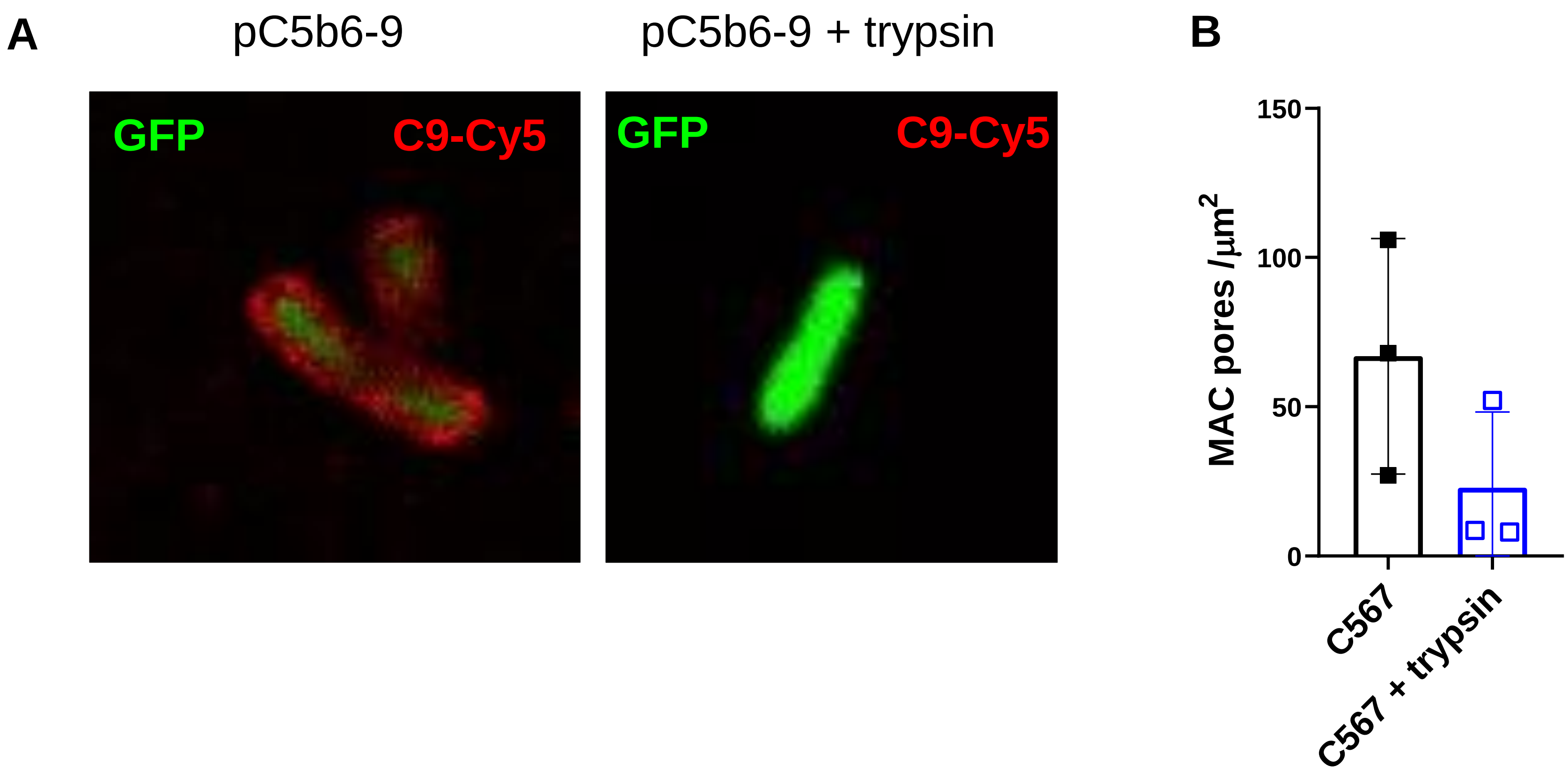
A
pC5b6-9
pC5b6-9 + trypsin
GFP
C9-Cy5
GFP
C9-Cy5
B
MAC pores /μm²
150
100
50
0
C567
C567 + trypsin
